# A multiplexed, target-based phenotypic screening platform using CRISPR interference in *Mycobacterium abscessus*

**DOI:** 10.1101/2025.03.17.643728

**Authors:** Donavan Marcus Neo, Ishay Ben-Zion, Josephine Bagnall, Matthew Solomon, Austin Bond, Emily Gath, Shuting Zhang, Noam Shoresh, James Gomez, Deborah T Hung

## Abstract

The rise of difficult-to-treat *Mycobacterium abscessus* infections presents a growing clinical challenge due to the immense arsenal of intrinsic, inducible and acquired antibiotic resistance mechanisms that render many existing antibiotics ineffective against this pathogen. Moreover, the limited success in discovery of novel compounds that inhibit novel pathways underscores the need for innovative drug discovery strategies. Here, we report a strategic advancement in PROSPECT (PRimary screening Of Strains to Prioritize Expanded Chemistry and Targets), which is an antimicrobial discovery strategy that measures chemical-genetic interactions between small molecules and a pool of bacterial mutants, each depleted of a different essential protein target, to identify whole-cell active compounds with high sensitivity. Applying this modified strategy to *M. abscessus*, in contrast to previously described versions of PROSPECT which utilized protein degradation or promoter replacement strategies for generating engineered hypomorphic strains, here we leveraged CRISPR interference (CRISPRi) to more efficiently generate mutants each depleted of a different essential gene involved in cell wall synthesis or located at the bacterial surface. We applied this platform to perform a pooled PROSPECT pilot screen of a library of 809 compounds using CRISPRi guides as mutant barcodes. We identified a range of active hits, including compounds targeting InhA, a well-known mycobacterial target but under-explored in the *M. abscessus* space. The unexpected susceptibility to isoniazid, traditionally considered to be ineffective in *M. abscessus*, suggested a complex interplay of several intrinsic resistance mechanisms. While further complementary efforts will be needed to change the landscape of therapeutic options for *M. abscessus*, we propose that PROSPECT with CRISPRi engineering provides an increasingly accessible, high-throughput target-based phenotypic screening platform and thus represents an important step towards accelerating early-stage drug discovery.

## INTRODUCTION

The genus *Mycobacterium* comprises of almost 200 diverse environmental and pathogenic species, most notably divided into slow-growing species like *Mycobacterium tuberculosis*, and fast-growing ones like *Mycobacterium smegmatis*.^1–3^ Apart from the well-known disease causing species including *Mycobacterium tuberculosis* and *Mycobacterium leprae,* which cause tuberculosis and leprosy, respectively, other species are collectively referred to as nontuberculous mycobacteria (NTMs). Over the past few decades, incidence of NTM infections have been on the rise globally despite limitations in diagnosis and reporting, causing infections that range from skin and soft tissue to pulmonary disease.^3–5^ While certain risk factors can predispose patients to NTM infections, such as chronic lung disease (*e.g.*, cystic fibrosis, chronic obstructive pulmonary disease), immunodeficiency states, and post-invasive procedures, infections have also been increasingly reported in otherwise healthy individuals.^3, 6–11^

One NTM in particular, *Mycobacterium abscessus,* has gained notoriety as a highly resistant organism leading to difficult-to-treat infections.^6–9, 11^ Treatment regimens are largely empirical and involve multiple drugs.^12–15^ Nonetheless, treatment success rates of *M. abscessus* remain unsatisfactorily low due to its large arsenal of intrinsic, inducible and acquired antibiotic resistance mechanisms.^6–9, 11–19^ As a result, many currently available antibiotics used to treat other infections are rendered ineffective against *M. abscessus*, including common anti-tuberculosis agents such as isoniazid and rifampicin. Taken together, its unsurprising *M. abscessus* has earned the moniker of being an “incurable nightmare”, underscoring its clinical relevance as an emerging pathogen with a clear unmet clinical need.

Exacerbating the lack of effective treatment options, the drug discovery pipeline remains surprisingly scarce.^18, 20–23^ Extremely low hit rates have rendered conventional whole-cell screening assays largely unsuccessful in identifying candidate molecules while biochemical target-based approaches can produce inhibitors with high target affinity but lack whole-cell activity.^24^.^18, 21, 25–27^ Consequently, drug development for *M. abscessus* has largely focused on repurposing approved antibacterials (*e.g.*, clofazimine, bedaquiline), repurposing lead candidates against a limited set of targets (*e.g.*, inhibitors against penicillin-binding proteins, lipid transporter Mycobacterial membrane protein Large 3 MmpL3 and DNA gyrase),^18, 20–23^ and identifying compounds/combinations that can circumvent intrinsic resistance mechanisms (*e.g.*, rifabutin, β-lactamase combinations).^28–34^ Clearly, new screening strategies are needed that can improve discovery of novel compounds that inhibit novel targets. As such, strategies that could concurrently prioritize the discovery of potential cell-active hit compounds against *M. abscessus* while identifying putative targets/pathways would be ideal.

We recently published a method termed PROSPECT (PRimary screening Of Strains to Prioritize Expanded Chemistry and Targets) as a new strategy for antibiotic drug discovery. PROSPECT identifies new molecules that can be prioritized based on biological insight gained from the primary screening data and that ultimately can lead to the development of new antibiotic candidates that would have eluded conventional discovery.^24, 35–37^ The first genome-wide application of PROSPECT to *M. tuberculosis* involved multiplexed screening of a pool of barcoded, engineered mutant strains, each depleted of a different essential target (*i.e.* hypomorphs).^24, 36^ The hypersensitivity of some strains due to target depletion allowed the identification of active small molecules despite the absence of wild-type activity thereby increasing the numbers of active candidates that can be discovered, with subsequent chemical optimization to achieve wild-type activity, while simultaneously associating the molecules with putative targets or mechanisms of action based on the specificity of hypersensitivity in some strains over others. In a second, more limited (mini) version of PROSPECT, we identified valuable probes against the gram negative pathogen Pseudomonas aeruginosa by screening a pool of mutants depleted for essential outer membrane targets.^37^

Here we now report adapting PROSPECT for *M. abscessus* with an important key modification to make the platform far more accessible. Essential gene knockdown in *M. tuberculosis* was achieved using a target proteolysis strategy or with an inducible promoter for transcriptional control,^36^ while a constitutive, promoter replacement strategy was used for *P. aeruginosa*.^37^ All of these strategies required laborious homologous recombination efforts.^24, 36, 37^ Here, we leveraged advances in CRISPR interference, harnessing a system where a dead Cas9 from *Streptococcus thermophilus* CRISPR1 (*Sth1* dCas9) locus can be easily programmed in mycobacteria to easily achieve transcriptional interference by varying the targeting sgRNA sequence.^38–40^ The CRISPR guides (sgRNA) themselves can simultaneously serve as barcodes to enable multiplexing of mutants in a pooled screen. Applying CRISPRi, we focused on a subset of essential genes (termed mini-PROSPECT)^37^ localized on the surface of *M. abscessus* or involved in cell wall synthesis, given the challenges of achieving intracellular small-molecule accumulation in this challenging pathogen. We thus report the development of a CRISPRi-based, mini-PROSPECT assay in *M. abscessus* targeting a pool of 60 engineered hypomorphs. We performed a pilot screen of a library of 809 compounds with known antibiotic activity (Figure 1) as proof of principle and identified InhA as a potentially under-explored antibacterial target in *M. abscessus* by shedding light on its mechanisms of intrinsic resistance to the well-known InhA inhibitor, isoniazid.

**Figure 1:**
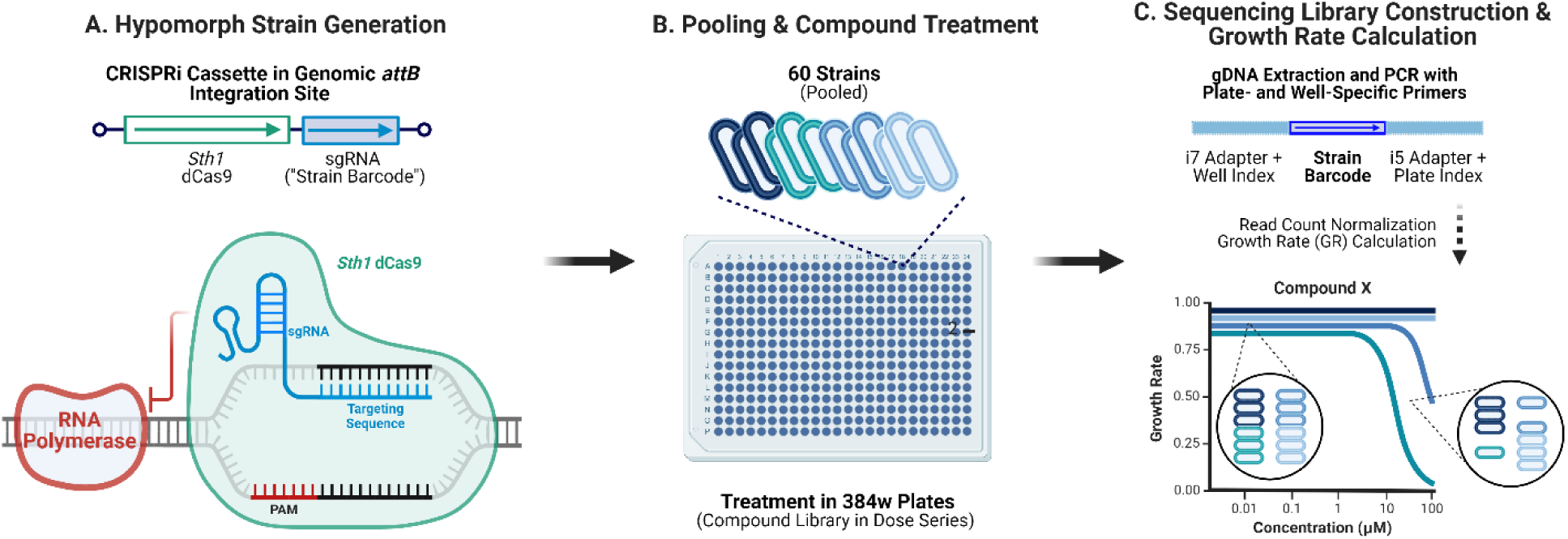
Overview of the multiplexed, CRISPRi-based, min-PROSPECT assay in *M. abscessus*. **A.** CRISPRi serves as a versatile tool for easily generating hypomorphs through transcriptional control by varying the sgRNA targeting sequence. The sgRNA sequence is unique for each strain and concurrently serves as a strain barcode. **B.** A pilot screen was performed pooling 60 hypomorph strains against a library of 809 compounds with known antibacterial activity in 384-well format in 8-point dose-response. **C.** By amplifying the strain barcode (sgRNA guide sequence) with plate– and well-indices, we apply a sequencing-based read-out of strain growth to measure the census of each mutant in response to each small-molecule perturbation, thereby generating chemical-genetic interaction profiles that shed light on putative targets or mechanisms of action. Created with Biorender.

## RESULTS

### Identifying essential genes in M. abscessus using Tn-Seq and FiTnEss

We performed a genome-wide negative selection study to define the essential targets localized on the surface of *M. abscessus* or involved in cell wall synthesis. We performed Himar1 Tn-seq in the reference strain *M. abscessus* ATCC 19977 using the established mycobacteriophage ΦMycoMarT7 to first identify all essential genes.^41–43^ We achieved ∼10^7^ mapped reads per library and high coverage of TA sites (> 70%), reads at each TA site were concordant between duplicate libraries (R^2^ = 0.8954), and reads per gene formed a characteristic bimodal distribution (Supplementary Figure 1). Using the FiTnEss analytical algorithm,^44, 45^ we identified 402 essential genes (ES), 4203 non-essential genes (NE), and 223 genes with intermediate classification we defined as “growth defective” (GD) (Supplementary Data). 92 genes lacked usable TA sites for the FiTnEss pipeline. We further analyzed the Tn-seq datasets using the commonly applied Hidden Markov Model (HMM) and found high concordance in essentiality calls (93.7%) between the two methods (Supplementary Table 1, Supplementary Data).^45–47^ Moreover, HMM analysis had high agreement with those from recently published works defining essential genes in *M. abscessus*.^48, 49^ Based on the Tn-Seq results, we identified 28 target genes (Table 1) for hypomorph generation, largely focusing on essential processes at the outer membranes such as cell wall synthesis,^50–53^ with some additional genes that encode known, validated drug targets in *M. tuberculosis*.^53–55^

**Table 1:** List of target genes selected for hypomorph generation.

| **Gene** | **Locus** | **FiTnEss\*** | **HMM\*** | **General Function** |
| --- | --- | --- | --- | --- |
| *rpoB* | MAB_3869c | ES | ES | DNA Transcription |
| *gyrB* | MAB_0006 | ES | ES | DNA Synthesis |
| *gyrA* | MAB_0019 | GD | ES | DNA Synthesis |
| *kasA* | MAB_1877c | ES | ES | Mycolic Acid Biosynthesis |
| *inhA* | MAB_2722c | ES | ES | Mycolic Acid Biosynthesis |
| *fadD32* | MAB_0179 | ES | ES | Mycolic Acid Biosynthesis |
| *mmpL3* | MAB_4508 | GD | ES | Mycolic Acid Biosynthesis |
| *dprE1* | MAB_0192c | NE | GD | Arabinogalactan Biosynthesis |
| *glf* | MAB_0170 | ES | ES | Arabinogalactan Biosynthesis |
| *rfbE* | MAB_0203c | ES | GD | Arabinogalactan Biosynthesis |
| *aftA* | MAB_0190c | ES | ES | Arabinogalactan Biosynthesis |
| *aftB* | MAB_0174 | ES | GD | Arabinogalactan Biosynthesis |
| *aftC* | MAB_2978 | GD | ES | Arabinogalactan Biosynthesis |
| *aftD* | MAB_4472 | ES | ES | Arabinogalactan Biosynthesis |
| *murJ* | MAB_4937 | NE | ES | Peptidoglycan Biosynthesis |
| *pbpB* | MAB_2000 | ES | ES | Peptidoglycan Biosynthesis |
| *lpqW* | MAB_1315 | ES | ES | Glycolipid Biosynthesis |
| *mptA* | MAB_1991c | ES | ES | Glycolipid Biosynthesis |
| *qcrB* | MAB_1966c | GD | GD | ATP Synthesis |
| *atpB* | MAB_1447 | ES | ES | ATP Synthesis |
| *ndh* | MAB_2429c | ES | ES | ATP Synthesis |
| *sdhC* | MAB_3673 | ES | ES | ATP Synthesis |
| *sdhA* | MAB_3675 | ES | ES | ATP Synthesis |
| *sdhB* | MAB_3676 | ES | ES | ATP Synthesis |
| *secY* | MAB_3784c | ES | ES | Protein Secretion |
| *dsbA* | MAB_3269c | ES | GD | Disulfide Bond Formation |
| *vkor* | MAB_3268c | ES | GD | Disulfide Bond Formation |
| *pbp-lipo* | MAB_3167c | ES | ES | Cell Morphology & Division |

\* Tn-seq essentiality calls abbreviated as ES for essential, GD for growth defect or NE for non-essential.

### Rapid generation of hypomorphic M. abscessus strains using CRISPRi

We chose a CRISPRi strategy to engineer *M. abscesses* mutants depleted for the 28 essential targets that could be used in mini-PROSPECT. To increase the efficiency of strain construction, given the number of strains we wished to construct, we adapted the methods previously reported in *M. abscessus* using the CRISPRi plasmid pJR965 for generating single gene depletions.^49, 56–58^ In contrast to these one-step methods, which have high rates of background resistance to the antibiotic selection marker (kanamycin), we utilized a two-step method (Supplementary Figure 2) analogous to one we recently published for CRISPR genome editing in *M. abscessus*.^59^ In brief, we first introduced into *M. abscessus* a plasmid containing a fluorescent mCherry reporter under the control of an anhydrotetracycline-inducible reporter but lacking any tetracycline repressor (TetR). Next, transforming this parental mCherry-expressing strain with the CRISPRi plasmid pJR965 introduces the TetR gene which encodes the repressor and suppresses mCherry expression in the absence of anhydrotetracycline (AHT). This workflow allowed us to accurately distinguish correct kanamycin-resistant transformants that have turned white from background mutants that remained pink but acquired spontaneous antibiotic resistance during selection (Supplementary Figure 3). Through this two-step method, we were able to quickly generate large numbers of hypomorphic strains.

For each of the 28 target genes, we designed 2-3 sgRNAs with moderate to strong protospacer adjacent motifs (PAMs) using previously established methods,^38–40^ and introduced them into *M. abscessus.* CRISPRi induction with AHT in each hypomorph led to varying levels of growth defect when measured in single-plex using standard OD_600_ measurements (Figure 2A). An additional four strains were also engineered with non-targeting control sgRNAs as surrogate wild-type controls (WT), with no growth defect observed with CRISPRi induction.

**Figure 2:**
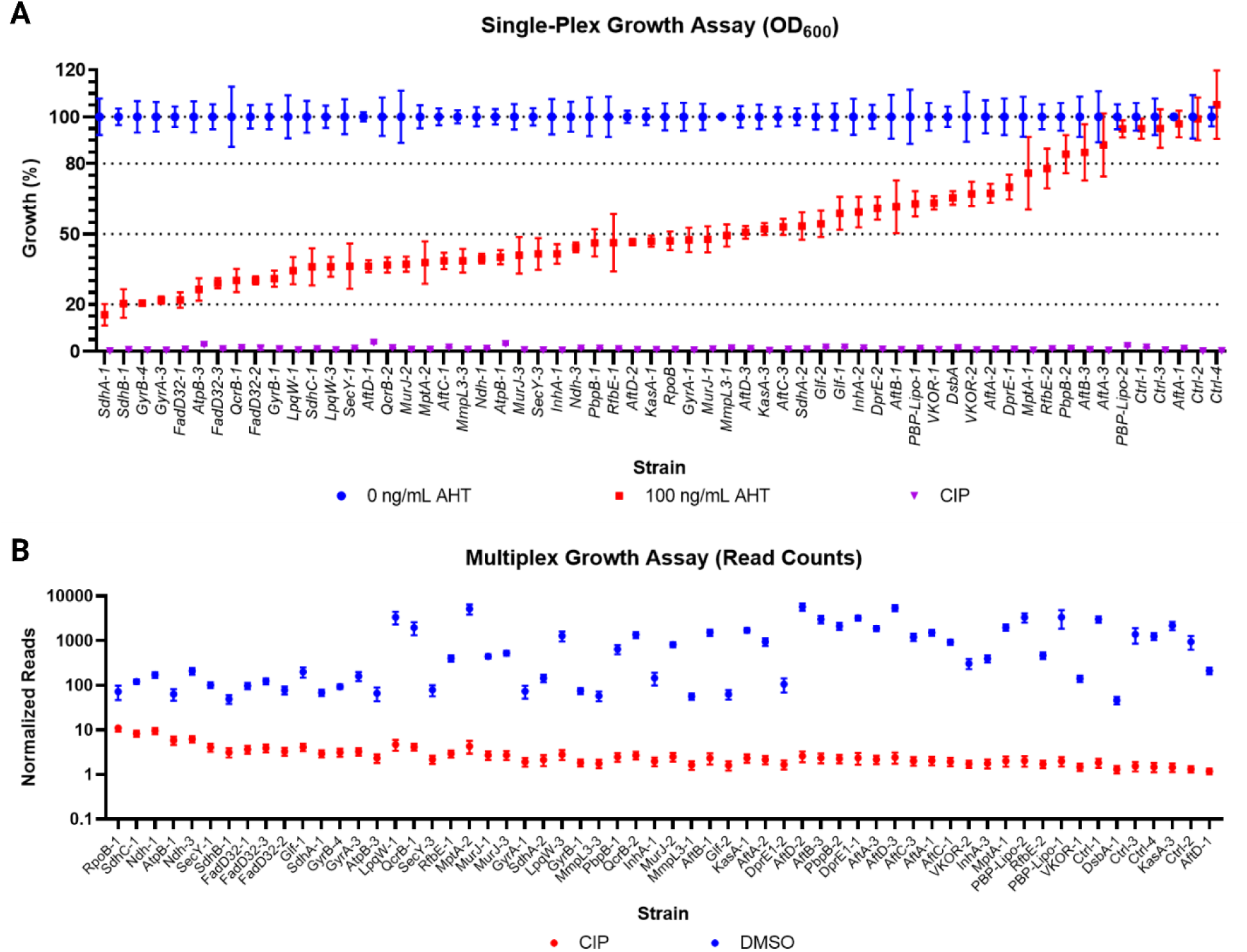
Growth of the 60 hypomorph strains generated in single-plex and multiplex. **A.** Strain growth rates measured in single-plex. Growth (%) was calculated based on OD_600_ values in liquid medium normalized to growth under uninduced CRISPRi conditions after 4 days (blue). Growth defects were seen in a majority of hypomorphs when CRISPRi was induced with 100 ng/mL anhydrotetracycline (AHT) (red). Ciprofloxacin (CIP) was used as a positive well-killing control (purple). Data are mean values and standard deviations from four technical replicates. **B.** Growth assay in multiplex. Strain abundance was measured using read counts as a surrogate after normalization to account for any variation in DNA extraction and PCR amplification between and within plates. CIP was used as a positive well-killing control (red) while DMSO was used as a negative control (blue). Median normalized read counts were obtained from twelve wells per plate. Data represents the mean values and SEM from seventy-two technical replicate plates.

### Adapting the CRISPRi-based PROSPECT assay to M. abscessus

We utilized a similar sequencing-based strategy for mini-PROSPECT in *M. abscessus* as previously reported, using sequencing reads of individual mutant barcodes as a means to enumerate strain census within a pool as a consequence of small molecule exposure (Figure 1).^24, 36^ In this case, we took advantage of the CRISPRi guide as the barcode uniquely identifying each mutant. Strain abundance within the pool was measured after 4 days of compound exposure by extracting crude genomic DNA, amplifying the barcode with plate and well indices, and then sequencing of barcode amplicons (sgRNA sequences) to enumerate strain census in each well. Reads were normalized to account for any variation in DNA extraction and PCR amplification between and within assay plates.

Compared to PROSPECT for *M. tuberculosis* where only one hypomorph strain per essential gene was included,^24^ we included multiple *M. abscessus* hypomorphs per gene spanning a range of growth rates. To account for these growth differences, we constructed the hypomorph pool with the four WT control strains at a relative abundance of 1× each, and with the hypomorphs at relative abundances from 1× to 120×, depending on their baseline growth rates. This was a necessary adjustment to constrain endpoint read counts across all strains to within two to three orders of magnitude to achieve adequate sequencing depth for each strain (Figure 2B).

### Pilot multiplexed screening against a library of known antibiotics

We performed a pilot screen using these 60 engineered *M. abscessus* strains against a small library of 809 compounds in an 8-point dose series, expanding on a set of 437 compounds we had previously assembled with annotated MOAs and known or predicted anti-tubercular activity, from strong mechanistic validation of MOAs to in silico protein docking.^60^ The screen thus corresponded to querying 22,652 unique compound-gene interactions. The screen was performed in duplicate and included DMSO as a negative control and ciprofloxacin as positive control.

Traditional metrics to quantify drug sensitization are often based on the absolute difference between treated and untreated growth rates.^24, 61^ These absolute metrics would thus have more extreme values for strains with faster baseline (untreated) growth rate. Since strains in our screen vary greatly in their baseline growth rates, this could lead to bias and prioritization of compounds that sensitize fast growing strains. We therefore followed our recent approach,^60^ and used the normalized Growth Rate (GR) metric that alleviates the bias by normalizing the treatment growth rate by the baseline (DMSO) growth rate (Methods).^61^ GR varies between 0 (representing full growth inhibition as in ciprofloxacin (CIP)-treated wells as a positive control) and 1 (representing no growth inhibition as in DMSO-treated wells) (Supplementary Figure 4A) and is independent of each strains’ baseline (DMSO) growth rate. To quantify how extreme a GR value is compared to the distribution of DMSO GR values, we standardized (z-scored) GR for a given strain at a given treatment to the DMSO GR values of that strain, yielding DMSO-Standardized GR (DSGR) (Methods), with DMSO-treated wells centering around DSGR = 0 and CIP-treated wells having negative DSGR values generally less than –5 (Supplementary Figure 4B). For most strains, *Z*’ factors > 0.5 (37 out of 60; Supplementary Figure 4C). For the remaining 23 strains, 20 had *Z’* factors between 0.25 and 0.5.

To define “hits”, we defined a metric *IC*^*GR*^_50_ as the concentration required to inhibit GR by 50% (*i.e.*, the lowest concentration for which GR value is below 0.5). Balancing activity and specificity, we then defined hits as any compound that has:

(1) an *IC*^*GR*^_50_ value in a hypomorph strain that is lower than the average *IC*^*GR*^_50_ across the 4 WT strains – meaning the hypomorph strain is hypersensitive to the hit compound (*i.e.*, a compound-gene interaction) relative to wild-type bacteria.
(2) a corresponding average DSGR value < –5 at 1× and 2× *IC*^*GR*^_50_ for that hypomorph strain – meaning the compound-gene interaction is significant. If the *IC*^*GR*^_50_ was the maximum concentration tested, then only the DSGR value at 1× *IC*^*GR*^_50_ was used.
(3) ≤ 12 hypomorph strains that fulfill criteria (1) and (2).

We identified 107 hits from 809 compounds screened (13.2%) (Table 2, Supplementary Data). These hits corresponded to 327 significant compound-gene interactions associated with compound hypersensitivity (1.44% of 22,652 possible pairs), with an average of ∼3 strains sensitized per hit compound. Breaking down these into their annotated general mechanism of action, 29 of 107 hits (27%) are annotated to affect cell shape and integrity, while 8 (7%) are annotated to affect respiration. Bias toward compounds with these mechanisms of action is consistent with the engineered hypomorph pool focusing on essential genes functioning at the membrane interface such as cell wall biosynthesis and ATP production. For instance, 52 out of the 63 significant compound-gene interactions associated with compounds annotated to affect cell shape and integrity involved genes responsible for biosynthesis of cell wall components (*e.g.*, mycolic acids, arabinogalactan, glycolipids *etc.*). Likewise, 21 out of the 28 significant compound-gene interactions associated with compounds annotated to affect respiration involved genes involved in the mycobacterial electron transport chain. Additionally, 41 hits (38%) are annotated to affect translation and replication, including known antibiotics commonly used against *M. abscessus* such as the macrolides and fluoroquinolones.

**Table 2:** Number of hits and their associated compound-gene pairs categorized by general mechanism of action.

| Annotated Mechanism of Action | Compounds | Compound-Gene Interactions |
| --- | --- | --- |
| Cell Shape/Integrity | 29 | 63 |
| Translation | 25 | 91 |
| Replication | 16 | 68 |
| Respiration | 8 | 28 |
| Others | 29 | 77 |
| **Total** | **107** | **327** |

To validate hits, we tested a set of these interactions using an orthogonal OD_600_-based single-plex growth assay. We cherry-picked 43 significant compound-gene interactions (representing 31 unique compounds) to validate, as well as 94 neutral compound-gene pairs (representing 68 unique compounds) that were not associated with any hypersensitivity as negative controls. This set of 137 compound-gene pairs represented 93 unique compounds. We similarly calculated GR and DSGR values using OD_600_ (Methods). To account for the general phenomenon of multiplexed assays being more sensitive to compound hypersensitization than single-plexed growth due to growth in competition, we lowered the threshold for calling significant interactions using a single-plex, OD_600_ based readout, as those with:

(1) an *IC*^*GR*^_20_ value in a hypomorph strain that is lower than that in the surrogate-WT strain, where *IC*^*GR*^_20_ is defined as the concentration required to inhibit GR by 20% (*i.e.*, concentration for a GR value of 0.8).
(2) a corresponding average DSGR value of < –4 at 1× and 2× *IC*^*GR*^_20_ for that hypomorph strain. If the *IC*^*GR*^_20_ was the maximum concentration tested, then only the DSGR value at 1× *IC*^*GR*^_20_ was used.

We confirmed 25 out of 43 significant interactions (positive predictive value = 58%), as well as 63 out of 94 negative control compound-gene interactions (negative predictive value = 67%). Taken together, the multiplexed pilot screen performed reasonably well and identified strong chemical-genetic interactions with an overall accuracy of 64% (Table 3).

**Table 3:** Confusion matrix of compound-gene pairs selected for validation.

| Significant chemical-genetic interaction? |  | Demultiplexed Assay |  | Overall Accuracy*<br>= 64.2% |
| --- | --- | --- | --- | --- |
|  |  | YES | NO |  |
| **Multiplexed Screen** | YES | 25 | 18 | PPV<sup>^</sup> = 58.1% |
|  | NO | 31 | 63 | NPV<sup>†</sup> = 67.0% |

\* Overall accuracy calculated as the number of concordant interactions (strong interaction or no interaction in both multiplexed and demultiplexed) divided by the total number tested.

### The intrinsic resistance of M. abscessus to isoniazid is multifactorial and mediated by multiple mechanisms

From the pilot multiplexed screen, 11 known *M. tuberculosis* InhA inhibitors, including isoniazid, had specific activity for the *M. abscessus inhA* hypomorph, out of 29 hits annotated to affect cell shape and integrity (Table 2). 9 of these hits were confirmed in demultiplexed testing against the *inhA* hypomorph. While the majority of validated demultiplexed interactions only showed shifts in *IC*^*GR*^_20_, three structurally related InhA inhibitors also showed lower MIC_90_ values in the *inhA* hypomorph compared to the control wild-type strain, suggesting a signficiant chemical-genetic interaction.

Surprisingly, isoniazid and 7 structurally related analogs had activity against the *inhA* hypomorph, despite isoniazid generally being considered inactive against *M. abscessus*. Isoniazid is a first-line anti-tubercular antibiotic which requires activation by the catalase-peroxidase KatG, followed by formation of a NAD^+^ adduct which then inhibits InhA, an essential enoyl acyl carrier protein reductase involved in mycolic acid biosynthesis.^62^ While isoniazid is very active in *M. tuberculosis*, its lack of activity against other mycobacterial species has been conventionally thought to be due to the inability of respective KatG orthologs to catalyze the activation of isoniazid.^63^ The activity of isoniazid against the *inhA* hypomorph suggested that the *M. abscessus* KatG ortholog must possess some degree of catalytic activity, at least enough to activate this class of compounds to yield the observed activity in the *inhA* hypomorph.

To understand the basis for isoniazid’s surprising activity in the *inhA* hypomorph, we overexpressed the *M. abscessus* and *M. tuberculosis* orthologs of *katG* in wild-type *M. abscessus* and the *inhA* hypomorph. Overexpression of *katG*_Mab_ in wild-type *M. abscessus* indeed slightly increased its susceptibility to isoniazid, thus confirming the ability of KatG_Mab_ to convert isoniazid to its active form. However, overexpression of *katG*_Mtb_ conferred even greater susceptibility both in wild-type *M. abcessus* and the *inhA* hypomorph, with the latter strain now having an MIC_90_ of 3.13 μg/mL (Figure 3A). (Comparable expression levels of both *katG orthologs* were confirmed by qRT-PCR (data not shown)). Thus, although KatG_Mab_ has less isoniazid-activating activity than KatG_Mtb_, it still possesses some capacity to activate isoniazid. Of note however, even when isoniazid can be optimally activated in *M. abscessus* by expression of the *M. tuberculosis katG*_Mtb_ ortholog, the MIC_90_ of isoniazid in remained over 60-fold higher than that in wild-type *M. tuberculosis* (MIC_90_ = 0.03-0.05 μg/mL), thus indicating that intrinsic resistance to isoniazid in *M. abscessus* is more complex than a lack of sufficient KatG-dependent activation.

**Figure 3:**
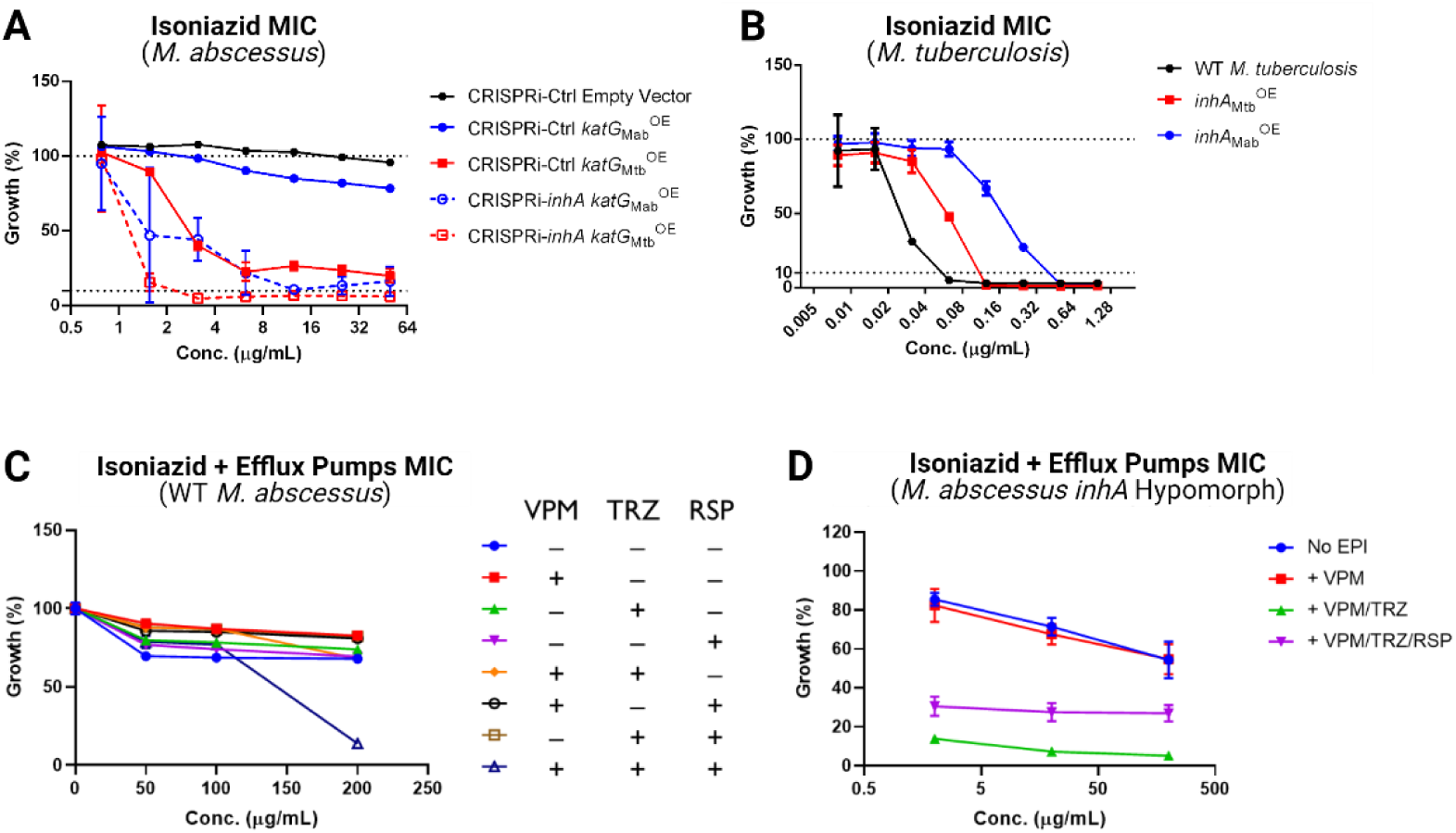
Isoniazid dose-response curves in *M. abscessus* and *M. tuberculosis* to unveil intrinsic isoniazid resistance mechanisms in *M. abscessus*. Growth (%) was calculated based on OD_600_ values in liquid medium normalized to DMSO-treated (A-C) or efflux pump inhibitor (EPI) only wells. Data are mean values and standard deviations from four technical replicates. **A.** *katG*_Mtb_ overexpression confers greater sensitivity to isoniazid than *katG*_Mab_ in CRISPRi-Ctrl WT-surrogate *M. abscessus* strain. Dose-response curves representing empty vector (black), *katG*_Mab_^OE^ (blue) or *katG*_Mtb_^OE^ (red) with either CRISPRi-Ctrl (solid lines) or CRISPRi-*inhA* knockdown (dotted lines). **B.** *inhA*_Mab_ overexpression confers greater resistance to isoniazid than *inhA*_Mtb_ overexpression in *M. tuberculosis*. Dose-response curves representing empty vector (black), *inhA*_Mab_^OE^ (blue) or *inhA*_Mtb_^OE^ (red) in *M. tuberculosis*. **C.** Only co-treatment with three efflux pump inhibitors (EPI) improves activity of isoniazid against WT *M. abscessus.* EPIs were added simultaneously with an isoniazid dose-response series. EPIs were added in fixed concentrations as follows: verapamil (VPM) (125 μg/mL = 1/4 × MIC_90_), thioridazine (TRZ) (7.8 μg/mL = 1/4 × MIC_90_), reserpine (RSP) (12.5 μg/mL = 1/4 × maximum concentration tested). **D.** Co-treatment with both VPM and TRZ improves isoniazid activity in *M. abscessus inhA* hypomorph strain (CRISPRi-*inhA*). Hypomorph strain was treated with isoniazid without EPI (No EPI), with VPM alone (one EPI), with VPM and TRZ (two EPIs), or with VPM, TRZ and RSP (three EPIs) at the concentrations described above.

To investigate this further, we compared the transcriptional levels of *katG* and *inhA* in *M. abscessus* and *M. tuberculosis* to rule out differences in their expression levels as a contributing factor to their difference in susceptibility. However, there were no significant differences in either gene expression levels (data not shown). We then sought to investigate if there were any species-specific differences in *inhA* orthologs that could play a role in the differing susceptibilities. We episomally overexpressed the *M. abscessus* and *M. tuberculosis inhA* orthologs in wild-type *M. tuberculosis*. The *inhA*_Mab_ overexpressing strain had a 10-fold higher MIC in the *inhA*_Mab_ overexpressing strain (MIC_90_ = 0.395 μg/mL) compared to wild-type *M. tuberculosis* (MIC_90_ = 0.034 ug/mL) (Figure 3B), while overexpression of *inhA*_Mtb_ only led to 3-fold higher isoniazid MIC (MIC_90_ = 0.097 ug/mL). (There was only slightly higher expression levels of *inhA*_Mtb_ compared to *inhA*_Mab_, and thus cannot account for the observed differences (data not shown)). Therefore, the InhA_Mab_ ortholog is intrinsically more resistant to inhibition by isoniazid than the InhA_Mtb_ ortholog, thus also factoring into the relative resistance of *M. abscessus* to isoniazid.

Finally, analysis of the transcriptional response of *M. abscessus* to isoniazid also implicated induction of efflux activity as another contributing factor to *M. abscessus* relative resistance. Treatment of wild-type *M. abscessus* with isoniazid (250 μg/mL) resulted in significant differential expression of over 1000 genes by RNA-seq (Supplementary Figure 5A-C), including up-regulation of *inhA* and other genes in the mycolic acid biosynthesis pathway, the isoniazid-induced genes in the *iniBAC* operon,^64, 65^ and notably, over 20 genes from various efflux transporter families including major facilitator superfamily (MFS) transporters, ATP-binding cassette (ABC) transporters, and resistance-nodulation-cell division (RND) superfamily transporters, the latter including the mycobacterial membrane protein (MmpS/L) transporters.^66–68^ Of note, while some efflux pumps have been implicated as a cause of isoniazid resistance in *M. tuberculosis*,^69–71^ several putative efflux pumps upregulated in *M. abscessus* during isoniazid treatment such as the MFS pumps MAB_2263c and MAB_0069, and the amino acid-metabolite transporters MAB_0677c and MAB_3369 have no known orthologs in *M. tuberculosis*. Notable changes in selected genes were confirmed by qRT-PCR (Supplementary Figure 5D).^72, 73^ To test the role of inducible efflux activity contributing to intrinsic resistance in *M. abscessus*, we co-treated wild-type *M. abscessus* with known efflux pump inhibitors. While treatment with individual efflux pumps inhibitors verapamil,^17, 74–76^ thioridazine^17^ and reserpine^76^ had no impact on isoniazid efficacy when treated in single or two-drug combinations, treatment with all three efflux pump inhibitors did lower the MIC_90_ of isoniazid in wild-type *M. abscessus* (Figure 3C). Meanwhile, in the *inhA* hypomorph, the combination of verapamil and thioridazine (with or without reserpine) improved the activity of isoniazid (Figure 3D). Thus, the complex array of efflux systems also plays a role in the inferior activity of isoniazid for *M. abscessus*.

Taken together, contrary to previous assumptions that *M. abscessus* has an inactive KatG ortholog which renders isoniazid inactive, the finding that isoniazid is active against a *inhA* hypomorph in mini-PROSPECT screening has revealed a more complex interplay of factors that contribute towards intrinsic resistance to isoniazid. These factors include a less catalytically active KatG ortholog, intrinsic resistance of the InhA ortholog to inhibition by the activated isoniazid-NAD adduct, and a complex array of efflux pumps which likely work in tandem to reduce isoniazid accumulation within *M. abscessus*.

## DISCUSSION

We have developed a method PROSPECT that combines target-based and phenotypic screening to identify novel compounds that kill bacteria. By screening a pool of hypomorphs each depleted for a different essential target, PROSPECT can identify compounds that work against a broad range of targets and provide biological insight from the primary screen enabling hit prioritization based not solely on chemical structure and potency, but also specific biological activity.^24, 35–37^ Importantly, because activity is identified against hypomorphic mutants, many which are hypersensitized to respective inhibitors, the assay is much more sensitive compared to screening wild-type bacteria, thereby the enabling the discovery of active compounds, even if only active against hypomorphic strains, which can serve as starting points for chemical optimization to gain wild-type activity.^24, 35, 36, 60^ Given the dearth of molecules with any activity against wild-type *M. abscessus,* this could be a vital strategy for antibiotic discovery against a pathogen such as *M. abscessus*.

We have now advanced this strategy by applying CRISPRi technology to more easily engineer hypomorphic mutants depleted for specific essential targets, while leveraging the CRISPRi guides as barcodes to enable multiplexed screening (Figure 1).^38–40^ Previously, CRISPRi has been used to perform pooled, genome-wide screens *M. tuberculosis*, to investigate chemical-genetic interactions^40, 77, 78^ with a small number of known antibiotics. Here, with the goal of doing large-scale chemical screens to identify new, active compounds against *M. abscessus,* we adapted previous methods for constructing CRISPRi strains in *M. abscessus* to enable greater efficiency in generating hypomorphic strains for pooled screening (Supplementary Figure 2).^59^ By dramatically simplifying strain construction and the assay, with introduction of sgRNA sequences being far simpler than modifying the native copy of each gene with an additional step of introducing a barcode, the PROSPECT strategy can be more easily and broadly applied to many species of interest including the difficult-to-treat pathogen *M. abscessus*. Further, given the ease of strain construction and the smaller target set, we were able to include 2-3 hypomorph strains per target gene in the pool, each with slightly different levels of knockdown including strains with significant growth defects that still performed adequately (*Z*’ factors ≥ 0.4-0.5). Inclusion of these strains with more significant target depletion and thus growth defect could increase the chances of observing hypersensitization to inhibitors. This is in contrast to the original PROSPECT screen in *M. tuberculosis*, where only a single strain per gene possessing near normal growth relative to wild-type was included to ensure good assay performance; this restriction potentially came at the theoretical cost of using strains wherein the level of knockdown might not have been sufficient to result in hypersensitization to small molecule inhibitors.^24, 36^

The increased sensitivity of the mini-PROSPECT screen is evident in the 107 hit compounds identified, with 64% accuracy. Amongst these hits, only 30 of them had activity against wild-type *M. abscessus* (*IC*^*GR*^_50_ in surrogate-WT strain ≤ maximum concentration tested), and the additional 77 compounds would not have been identified solely based wild-type activity. Thus, strategies such as PROSPECT could be vital to identifying more active compounds against this extremely challenging bacterial species that could be starting points for further development, even if the original hit has no or limited initial wild-type activity. In the case of *M. tuberculosis,* we have demonstrated the ability of identifying initial hits with no such wild-type activity and the subsequent ability to achieve potent wild-type activity through medicinal chemistry efforts given an active starting scaffold.

The principle that PROSPECT is more sensitive for identifying inhibitors, even if they lack potent wild-type activity is evidenced by the identification of several significant interactions between the *inhA* hypomorphs and known InhA inhibitors, including the well-known antitubercular agent isoniazid, despite most of them having no wild-type activity. This was surprising given that isoniazid is expected to have limited activity against other mycobacterial due to the lack of a catalytically active KatG ortholog to activate isoniazid (and other KatG-dependent inhibitors).^62, 63^

Here we showed that the intrinsic resistance of *M. abscessus* for isoniazid is multifactorial, including the diminished albeit demonstrable ability of *M. abscessus* KatG relative to *M. tuberculosis* KatG to activate isoniazid, less effective inhibition or target engagement of *M. abscessus* InhA by the isoniazid-NAD adduct compared to *M. tuberculosis* inhA, and *M. abcessus’s* formidable array of redundant efflux pumps that can efficiently minimize intracellular drug concentrations. Interestingly, genetic comparisons of *katG* and *inhA* genes from *M. tuberculosis* and *M. abscessus* demonstrate their remarkable similarity. *M. abscessus* KatG has relatively high amino acid identity (72%) and similarity (83%) to the *M. tuberculosis* ortholog and most common mutated amino acid residues that confer isoniazid resistance in *M. tuberculosis* are not found in the *M. abscessus* ortholog.^79^ Meanwhile, the *M. abscessus* InhA ortholog shares 89% amino acid sequence identity (96% similarity) with the *M. tuberculosis* ortholog. While mutations within the coding region of *inhA* are only infrequently observed to confer resistance in *M. tuberculosis*,^80^ these are also not found in the *M. abscessus* ortholog. These small differences, along with a panoply of additional efflux pumps, result in two mycobacterial species that are extremely divergent in their isoniazid susceptibilities.

In summary, we have demonstrated for the first time the application of CRISPRi in *M. abscessus* for the purposes of multiplexed screening to investigate chemical-genetic interactions in high throughput for the purposes of compound discovery. In doing so, it dramatically facilitates the implementation of PROSPECT, thus enabling the expansion of chemical and target space for antibiotic discovery for critical, recalcitrant pathogens such as *M. abscessus.* While clearly additional parallel, complementary efforts will be required to truly change the landscape of treatment options for *M. abcessus,* including, for example, chemistry to achieve potent wild-type activity in candidates identified by PROSPECT and elucidation of physicochemical properties of small molecules that elude their complex efflux pump systems, we propose that CRISPRi-based PROSPECT could play a valuable role in early discovery efforts. While this proof-of-concept study focused on targets at the membrane interface, it is easy to envision expanding the hypomorph library to target more comprehensively all essential genes thereby leveraging the large number of potential, novel drug targets but which to date, lack known inhibitors, with the goal of ultimately expanding the numbers of possible candidate molecules with activity against *M. abscessus.* Even more broadly, combining CRISPRi engineering with a PROSPECT strategy can now be readily applied to numerous other clinically relevant bacterial species for which an antibiotic development pipeline is needed in the face of rising antibiotic resistance.

## MATERIALS AND METHODS

### Bacterial Strains and Growth Conditions

*Mycobacterium abscessus* ATCC 19977 (WT) was used as the parental wild-type strain for all experiments. *M. abscessus* was grown in Middlebrook 7H9 broth (BD) (M7H9) or Middlebrook 7H10 agar (BD) (M7H10) supplemented with 10% Middlebrook OADC (Oleic Albumin Dextrose Catalase) growth supplement (VWR), 0.5% glycerol (VWR) and 0.05% Tween-80 (Alfa Aesar) at 37°C. Transposon-sequencing experiments were conducted with growth media supplemented with 50 μg/mL (liquid) or 500 μg/mL (solid) kanamycin sulfate (KAN) (Sigma Aldrich). For strains transformed with various pJR965 CRISPRi plasmid derivatives, growth media was supplemented with 50 μg/mL (liquid) or 500 μg/mL (solid) KAN. For strains transformed with pCHERRY3 and pDN-Cherry derivatives, growth media was supplemented with 0.5 mg/mL (liquid) or 1 mg/mL (solid) hygromycin B (HYG) (Life Technologies). For induction of CRISPRi, growth media was supplemented with 100 ng/mL (liquid) or 250 ng/mL (solid) anhydrotetracycline hydrochloride (AHT) (Sigma-Aldrich).

For plasmid amplification, 5-alpha competent *Escherichia coli* (New England Biolabs) was used and grown in LB broth or agar supplemented with 50 μg/mL KAN or 150 μg/mL HYG as required.

For propagation and titration of mycobacteriophage ΦMycoMarT7, *Mycobacterium smegmatis* mc^2^155 was used and grown in M7H9 broth or M7H10 agar supplemented with 10% OADC and 0.5% glycerol (with or without Tween-80), according to previously reported methods.^81, 82^

For experiments in *Mycobacterium tuberculosis*, wild-type H37Rv was used and grown in M7H9 supplemented with 10% OADC, 0.2% glycerol, and 0.05% Tween-80 at 37°C. For strains transformed with overexpression plasmids, growth media was supplemented with 50 μg/mL HYG.

### Generation of Transposon Libraries in M. abscessus

Phage stocks were propagated and titrated in *M. smegmatis* as previously reported.^81, 82^ For each *M. abscessus* transposon library, 50-100 mL of cultures were grown to an OD_600_ of 1. Cultures were washed twice in Tween-free M7H9 broth and resuspended in MP buffer (50 mM Tris-HCl pH 7.5, 150 mM NaCl, 10 mM MgSO_4_, 2 mM CaCl_2_). Phage stock (> 10^11^ pfu/mL) was then incubated with resuspended cultures at a multiplicity of infection (MOI) of 7.5 for 6-8 hours at 37°C with shaking. The infected cultures were washed in phosphate-buffered saline (PBS) supplemented with 0.05% Tween-80 (PBS-Tween) and resuspended in 2 mL PBS-Tween. Serial dilutions were prepared in PBS-Tween to determine the titer while the remaining was plated on 245 mm square dishes containing M7H10 agar supplemented with 500 μg/mL KAN. After incubation for 7 days, the lawn of transduced colonies on the 245 mm plates were scraped and resuspended in 50 mL 7H9 broth supplemented with 20% glycerol and incubated at 4°C overnight with shaking. The resulting libraries were then aliquoted and stored at –80°C.

*Preparation of Transposon-sequencing (Tn-seq) Libraries, Mapping and Analysis* Sequencing libraries were prepared according to protocols adapted from other mycobacteria.^81, 82^ 1 mL aliquots of the frozen transposon libraries were first thawed and recovered in 40 mL M7H9 agar supplemented with 500 μg/mL KAN for 6-8 hours at 37°C with shaking. The cultures were then diluted to an OD of 0.01 and incubated for 2 days at 37°C in triplicate cultures. Genomic DNA (gDNA) was extracted according to previously reported protocols.^82^ Sequencing libraries for Tn-seq were then subsequently prepared using a modified protocol based on previous work in *M. tuberculosis*.^81, 82^ Briefly, gDNA was sheared using NEBnext dsFragmentase (New England Biolabs) and purified with 2 × SPRI. End repair and A-tailing of sheared gDNA was performed with 2 × SPRI purification after each step, followed by adapter ligation with 1.5 × SPRI purification. Hemi-nested PCR amplification of the Tn-junctions was performed with 1.5 × SPRI purification after each PCR cycle. Amplicon sequencing was carried out at the Broad Institute Genomics Platform using Illumina HiSeq 2500 and sequencing read counts were mapped to TA sites on the reference genome as previously reported for *P. aeruginosa*.^44, 45^ Two independent transposon libraries were prepared for sequencing and analysis (Pearson correlation R^2^ = 0.8954)

For FiTnEss, the pipeline (https://github.com/broadinstitute/FiTnEss) was applied to the mapped reads and analyzed as previously described.^81, 82^ Two levels of stringency were used for making predictions: a maximal stringency adjustment using family-wise error rate (FWER) and a slightly more relaxed one using false discovery rate (FDR). Genes with an adjusted P value < 0.05 in both libraries with both FWER and FDR were considered true “essentials” (ES), while genes with adjusted P value < 0.05 in both replicates with FDR alone were assigned with “growth defect” (GD) calls. The remaining genes were considered non-essential (NE).

For the Hidden Markov Model (HMM), the established TRANSIT pipeline was used to determine essentiality categories.^46, 47^ Insertion counts from FiTnEss pipeline were further normalized using TTR normalization and averaged across replicates. In HMM, the essentiality of each individual TA site was determined based on local insertion density in the contiguous regions surrounding that site and the mean value of nonempty read counts at all TA sites.^46, 47^ Sites with near 0 counts were assigned as essential, sites with ∼ 1/10 mean nonempty read count value were assigned as growth defect, and all remaining sites were assigned as non-essential. Genes are then assigned based on the majority essentiality call of TA sites within the gene.

### Plasmid Construction

Primers used for the construction and verification of plasmids are listed in Supplementary Data. Plasmids generated through this work are listed in Supplementary Data. All enzymes used for cloning were purchased from New England Biolabs pJR965 (Addgene plasmid #115163; http://n2t.net/addgene:115163; RRID: Addgene_115163; gift from Jeremy Rock) was used for CRISPRi in *M. abscessus*.^38–40^ All targetable sites on the *M. abscessus* genome (NCBI Reference Sequence: NC_010397.1) were extracted based on the previously reported 15 protospacer adjacent motifs (PAMs) for Sth1 Cas9 (Supplementary Data: PAM Sites).^38–40^ Selected targeting guides were purchased as complementary oligos (IDT) and assembled into BsmBI-linearized pJR965 according to previously reported protocols.^38–40^ sgRNA sequences in this work are listed in Supplementary Data.

pCHERRY3 (Addgene plasmid #24659; http://n2t.net/addgene:24659; RRID: Addgene_24659; gift from Tanya Parish) was used as the template for the construction of the *katG* overexpression plasmids.^83^ To first create pDN-Cherry, the original mCherry promoter on pCHERRY3 was replaced with the constitutive strong mycobacterial promoter P_smyc_ amplified from pJR962 (Addgene plasmid #115162; http://n2t.net/addgene:115162; RRID: Addgene_115162; gift from Jeremy Rock). Next, P_smyc_ and the target gene for overexpression were amplified using NEBuilder HiFi DNA Assembly Cloning Kit (New England Biolabs) into pDN-Cherry. For *katG* overexpression in *M. abscessus*, the *katG* gene from *M. abscessus* (*katG*_Mab_, MAB_2470c) or *M. tuberculosis* (*katG*_Mtb_, Rv1908c) were amplified from gDNA extracted from respective species. For *inhA* overexpression in *M. tuberculosis*, the *inhA-hemH* operon from *M. abscessus* (*inhA*_Mab_, MAB_2721c-MAB_2722c) or *M. tuberculosis* (*inhA*_Mtb_, Rv1484-Rv1485) were amplified for cloning. Plasmids were transformed and amplified into 5-alpha competent *E. coli* (New England Biolabs), isolated with QIAprep Spin Miniprep Kit (Qiagen), and verified by sequencing prior to further cloning or transformation.

### Generation of M. abscessus CRISPRi Strains

Electroporation of *M. abscessus* strains was performed according to previously published protocols using the BioRad Gene Pulser Xcell (at 2500 V, 1000 Ω, and 25 μF), recovered in M7H9 broth for 6-8 hours and plated on M7H10 agar plates supplemented with antibiotics as required for 5-7 days. Bacteria strains generated in this work are listed in Supplementary Data.

For CRISPRi strains used in the multiplexed screening and demultiplexed validation, a two-step transformation workflow was performed with WT *M. abscessus* (Supplementary Figure 2). First, pCHERRY3 was transformed into WT strain and colonies were selected from M7H10 plates containing HYG after 5 days and grown in M7H9 with HYG for 2 days. Successful transformants that had taken up pCHERRY3 were identified visually as pink colonies. Next, this parent pCHERRY3-containing strain was transformed with pJR965 derivatives containing various sgRNA sequences. Colonies were selected from M7H10 plates with KAN and HYG after 5 days. Successful transformants that had taken up pJR965 were identified visually as white colonies and grown in M7H9 with KAN and HYG for 2 days. 5 μL of each culture was mixed vigorously with 100 μL of UltraPure™ DNase/RNase-free distilled water (ThermoFisher Scientific) before incubation at 100°C for 6 min. After cooling to room temperature, 5 μL of this heat-killed suspension was used as template for colony polymerase chain reaction (PCR). Primers targeting the integration sites (attL and attR) as well as a region within the Sth1 Cas9 cassette were used to verify successful integration of pJR965 (Supplementary Data). PCR products were separated and visualized on 2% (w/v) agarose gel.

### Generation of Overexpression Strains

For *katG* overexpression in *M. abscessus*, an alternate workflow was employed from the one described above for generating hypomorphs. Briefly, WT *M. abscessus* was first transformed with pJR965 cloned with either non-targeting control sgRNA (equivalent to CRISPRi strain Ctrl-1) or inhA-targeting sgRNA (equivalent to CRISPRi strain InhA-1). Successful transformants were verified by colony PCR as above before transformation with either pDN-Cherry (empty vector) or pDN-Cherry-katG-Mab (*katG*_Mab_^OE^) or pDN-Cherry-katG-Mtb (*katG*_Mtb_^OE^). Successful transformants in this step were then identified visually as pink colonies. Overexpression was then verified by qRT-PCR.

For *inhA* overexpression in *M. tuberculosis*, the WT H37Rv strain was electroporated according to previously published protocols using the BioRad Gene Pulser Xcell, recovered in M7H9 broth for 20-24 hours and plated on M7H10 agar plates supplemented with antibiotics as required for 20 days. Overexpression was then verified by qRT-PCR.

### Demultiplexed Growth Assays for CRISPRi Knockdown

*M. abscessus* hypomorph strains were grown to OD_600_ ∼ 0.8 to 1.0 in media supplemented with KAN and HYG. For liquid growth assays, cultures were diluted to 96-well clear bottom plates (Corning) (final initial OD_600_ = 0.0025) in four technical replicates with or without AHT. 1% DMSO (Sigma-Aldrich) was used as the negative untreated control and 25 μg/mL ciprofloxacin hydrochloride (CIP) (MP Biomedicals) was used as the positive control. Plates were incubated at 37°C without shaking for approximately 4 days before OD_600_ measurements using a SpectraMax M5 plate reader (Molecular Dimensions). Growth (%) was calculated by normalizing OD_600_ from treatment wells with OD_600_ from DMSO control wells and plotted using GraphPad Prism 9.4.0.

For spotting assays, 10-fold dilutions of cultures were prepared (OD_600_ = 0.0025 to 2.5 × 10–6) and spotted onto M7H10 plates supplemented with KAN and HYG with or without AHT in 5 μL droplets.

### Dose-Response Growth Assays

As above, *M. abscessus* strains were grown to OD_600_ ∼ 0.8 to 1.0 in media supplemented with KAN and HYG as necessary and diluted to 96-well clear bottom plates (Corning) (final initial OD_600_ = 0.0025). 1% DMSO and 25 μg/mL CIP were used as negative and positive controls respectively. Plates were incubated at 37°C without shaking.

For demultiplexed validation assays, WT-surrogate control strain and hypomorph strains were grown in 100 ng/mL AHT, treated with compound dilution series, and incubated for 3 days before OD_600_ measurements and calculation of GR (see below).

For *katG* overexpression in *M. abscessus*, strains were treated with isoniazid dilution series and incubated for 3 days before OD_600_ measurements. For co-treatment with efflux pump inhibitors, strains were treated with isoniazid with or without efflux pump inhibitors (verapamil, thioridazine and/or reserpine) in dilution series and incubated for 4 days before OD_600_ measurements. Growth (%) was calculated by normalizing OD_600_ from treatment wells with OD_600_ from DMSO control wells and plotted using GraphPad Prism 9.4.0.

For *inhA* overexpression in *M. tuberculosis*, in media supplemented with HYG as necessary and diluted to 96-well clear bottom plates (Corning) (final initial OD_600_ = 0.0025). 1% DMSO and 10 μM rifampin were used as negative and positive controls respectively. Plates were incubated at 37°C without shaking. Strains were treated with isoniazid dilution series and incubated for 7 days before OD_600_ measurements. Growth (%) was calculated by normalizing OD_600_ from treatment wells with OD_600_ from DMSO control wells and plotted using GraphPad Prism 9.4.0.

### Multiplexed Screening and Sequencing

Library of known antibiotics was constructed as previously reported.^24, 60^ Compounds were arrayed onto 384-well plates in 0.2 μL aliquots in a two-fold dose-series by Broad Institute Compound Management. Each assay plate also contained wells for DMSO (negative control) and CIP (positive control). Strains were grown separately and subsequently pooled between 1 × and 120 × relative abundances according to Supplementary Data. The pooled strains were induced with 100 ng/mL AHT and added to each well in 40 uL aliquots (final pooled OD_600_ = 0.0025). Assay plates were incubated at 37°C without shaking for approximately 4 days before being heat-killed at 80°C for 2-3 h.

Genomic DNA (gDNA) extraction, PCR amplification and sequencing library construction was adapted from PROSPECT screening in *M. tuberculosis*.^24, 36^ Briefly, equal volumes of heat-killed culture and 20% DMSO were mixed and incubated at 95°C for 15 min. An additional pJR965-derivative plasmid was generated (targeting sequence = GCTAGATGACTGCAGGGACTC) and used as a lysis-control barcode and spiked into 20% DMSO prior to mixing for subsequent read count normalization. 2 μL of each well was used for PCR (20 cycles) using Q5 Polymerase (New England Biolabs) with 5’ primer overhangs to add plate– and well-identification barcodes and sequences for Illumina NGS. PCR products were then pooled, cleaned-up using 2 × AMPure XP beads (Beckmann). Sequencing was carried out at the Broad Institute Genomics Platform using Illumina HiSeq 2500. The Concensus2 Python script was then used to count the co-occurrence of each combination of the three barcodes corresponding to plate and well coordinates, and strain identity (https://github.com/broadinstitute/Concensus2). These raw counts were then annotated with compound information based on the inferred plate and well coordinates, and with strain identity based on the strain barcode (defined as 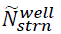 for a given well and strain).

### Computational Analysis of Sequencing Read Counts from Multiplexed Screening

To account for well-to-well differences in PCR efficiency and sequencing depth, we normalized raw counts for each well-strain pair (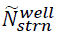) to the raw counts from the lysis-control spike-in for each well (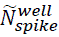). We first calculated the median log-transformed lysis-control spike-in counts across all wells using formula (1):

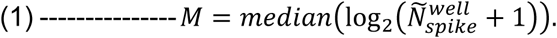

To avoid normalization of failed wells with very few reads, we removed wells with raw lysis-control counts lower than one eighth of the median (*i.e.*, 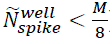). We then calculated normalized counts for each well-strain pair (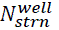) using (2):

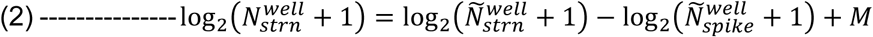

As the pooled strains exhibited variable baseline (DMSO) growth rates, we opted to use the normalized Growth Rate (GR) metric that is independent on baseline strain growth.^60, 61^ For each well-strain pair, we first calculated the estimated number of doublings (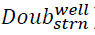) using the following formula (3):

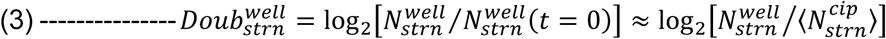

As time zero (*t* = 0) were not directly measured, we assumed 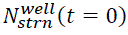 was proportional to the average read counts over all ciprofloxacin (CIP)-treated wells 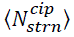, which is reasonable given no growth was observed in these wells and 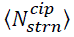 correlated well with relative initial strain abundances (Pearson correlation R^2^ = 0.8867).

Using average read counts over all DMSO-treated wells 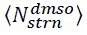 for each strain, we further calculated the estimated number of baseline doublings for each strain (4):

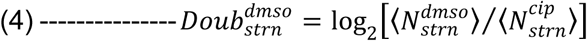

We then calculated normalized GR for each well-strain pair 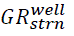 with formula (5), where GR varies between 0 (representing full growth inhibition as in CIP-treated wells) and 1 (representing no growth inhibition as in DMSO-treated wells). Negative GR values were transformed to zero as they were result of noise and do not have biological meaning.

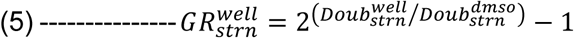

GR values used subsequently represent an average of two replicates per treatment condition (Pearson correlation R^2^ = 0.908, Supplementary Figure 4A). We further calculated *Z’* scores for each strain (as an estimate of strain performance using the formula (6):

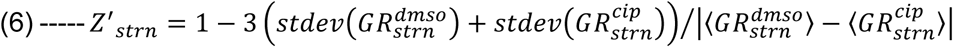

For each dose-response curve of GR against compound concentration, we defined a metric *IC*^*GR*^_50_ as the minimum concentration required to inhibit GR by 50% (*i.e.*, below a value of 0.5) if GR in the next dose (2 × *IC*^*GR*^_50_) is also less than 0.5. For inactive compounds where GR at the highest dose was > 0.5, we arbitrarily assigned *IC*^*GR*^_50_ as 2 × maximum concentration.

To quantify the significance of growth inhibition for each compound concentration-strain pair, we calculated the DMSO-standardized GR (DSGR) using formula (7), where median and median absolute deviation (MAD) were calculated across all DMSO-treatment wells.

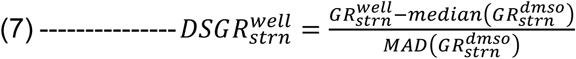

We then defined hits as any compound that has at least 1 hypomorph strain that has:

(1) an *IC*^*GR*^_50_ value in a hypomorph strain that is lower than the average *IC*^*GR*^_50_ across the 4 WT strains – meaning the hypomorph strain is hypersensitive to the hit compound (*i.e.*, a compound-gene interaction) relative to wild-type bacteria.
(2) a corresponding average DSGR value < –5 at 1× and 2× *IC*^*GR*^_50_ for that hypomorph strain – meaning the compound-gene interaction is significant. If the *IC*^*GR*^_50_ was the maximum concentration tested, then only the DSGR value at 1× *IC*^*GR*^_50_ was used.
(3) ≤ 12 hypomorph strains that fulfill criteria (1) and (2).

Processed multiplex screening read counts, GR and DSGR scores are available upon request.

### Validation of Hits using Demultiplexed Screening

Compounds were tested against individual strains in demultiplexed 96-well format according to methods described above. From blank-corrected OD_600_ values at day 3, we similarly calculated 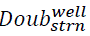 using OD_600_ values for CIP-treated wells as a surrogate for time zero OD_600_:

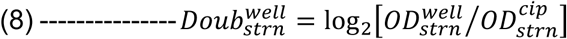

hen calculated GR (according to formula (5)). As we expected demultiplexed assays to be less sensitive than multiplexed screening, we defined *IC*^*GR*^_20_ is defined similarly as the minimum concentration required to inhibit GR by 20% (*i.e.*, concentration for a GR value of 0.8), and set lower thresholds for calling validated interactions as follows:

(1) an *IC*^*GR*^_20_ value in a hypomorph strain that is lower than that in the surrogate-WT strain, where *IC*^*GR*^_20_ is defined as the concentration required to inhibit GR by 20% (*i.e.*, concentration for a GR value of 0.8).
(2) a corresponding average DSGR value of < –4 at 1× and 2× *IC*^*GR*^_20_ for that hypomorph strain. If the *IC*^*GR*^_20_ was the maximum concentration tested, then only the DSGR value at 1× *IC*^*GR*^_20_ was used.

### RNA Extraction

For *M. abscessus*, strains were grown to OD_600_ ∼ 0.8 to 1.0 in M7H9 media containing antibiotics as required. Cultures were diluted to a final OD_600_ = 0.2 in 2.5 mL in 6-well plates (Corning). For *katG* overexpression, cultures were grown without addition of DMSO or any further compounds. For isoniazid-induced transcriptional changes, cultures were grown with either 1% DMSO or 250 μg/mL isoniazid. Plates were incubated at 37°C with shaking for approximately 6 hours before RNA extraction.

For *M. tuberculosis*, strains were grown to OD_600_ ∼ 0.4 to 0.8 in M7H9 media containing antibiotics as required before RNA extraction.

Briefly, 1.2-2.0 mL of each culture was spun down and resuspended in 0.5-0.7 mL TRIzol (Invitrogen) and stored at –80°C until ready for processing. Suspensions were transferred to FastPrep tubes (MP Biomedicals) containing ∼ 0.4-0.5 mL 0.1 mm zirconia/silica beads (BioSpec) and homogenized with a FastPrep bead beater (MP Biomedicals) at 10 m/s for 90 s. Samples were then incubated on ice for at least 3 minutes prior to addition of 200 μL 24:1 chloroform/isoamyl alcohol. Samples were mixed and centrifuged. The aqueous layer was then mixed with equal volume of 100% ethanol (Koptec) and processed with Direct-zol™ RNA MiniPrep kits (Zymo) with in-column DNase I treatment. RNA was eluted with DNase/RNase-free water, quantified using NanoDrop and stored in aliquots at –80°C.

### RNA Sequencing Library Construction and Analysis

Illumina cDNA libraries were generated using a modified version of the RNAtag-seq protocol.^84^ Briefly, 0.5 to 1 μg of total RNA was fragmented, depleted of genomic DNA, dephosphorylated, and ligated to DNA adapters carrying 5’-AN_8_-3’ barcodes of known sequence with a 5’ phosphate and a 3’ blocking group. Barcoded RNAs were pooled and depleted of rRNA using the RiboZero rRNA depletion kit (Epicentre). Pools of barcoded RNAs were converted to Illumina cDNA libraries in 2 main steps: (i) reverse transcription of the RNA using a primer designed to the constant region of the barcoded adaptor with addition of an adapter to the 3’ end of the cDNA by template switching using SMARTScribe (Clontech) as previously described;^85^ (ii) PCR amplification using primers whose 5’ ends target the constant regions of the 3’ or 5’ adaptors and whose 3’ ends contain the full Illumina P5 or P7 sequences. cDNA libraries were sequenced on the Illumina NextSeq 500 platform to generate paired end reads. Sequencing reads from each sample in a pool were demultiplexed based on their associated barcode sequence using custom scripts. Up to 1 mismatch in the barcode was allowed provided it did not make assignment of the read to a different barcode possible. Barcode sequences were removed from the first read as were terminal Gs from the second read that may have been added by SMARTScribe during template switching. Reads were then aligned *M. abscessus* ATCC 19977 genome (NCBI Reference Sequence: NC_010397.1) using BWA^86^ and read counts were assigned to genes and other genomic features using custom scripts. Differential expression analysis was conducted with DESeq2.^87^

### Quantitative RT-PCR (qRT-PCR)

Luna® Universal One-Step RT-qPCR kits (New England Biolabs) were used for all qRT-PCR experiments. qPCR primers were designed using Primer3 to amplify short amplicons (80-200 bp) and to have Tm of 60°C (± 0.5°C) and 40-60% GC. Primers used for qRT-PCR are listed in Supplementary Data. Fresh dilutions of RNA were prepared with final concentration of ∼ 5-10 ng/μL. Each reaction was prepared in triplicates according to established protocol. No-template control wells with no sample added, as well as No-RT control wells with no reverse transcriptase were also prepared. Relative expression was calculated using the ΔΔCt method. Briefly, average Ct values for each target gene was normalized to corresponding average Ct values for housekeeping genes in each sample to obtain ΔCt. 16S rRNA and *sigA* were selected as the housekeeping genes for *M. abscessus* and *M. tuberculosis*, respectively.

For verifying isoniazid-induced transcriptional changes in *M. abscessus*, ΔCt values for each sample was further normalized to the average ΔCt values for DMSO-treated samples for each target gene to obtain ΔΔCt. For verifying overexpression in *M. abscessus* and *M. tuberculosis*, ΔCt values for each sample was further normalized to either ΔCt in empty vector strain (*katG* overexpression), or to native copy of *inhA* (*inhA overexpression*) to obtain ΔΔCt. Relative expression or log_2_(fold-change) was defined as –ΔΔCt from three biologically independent replicates and plotted using GraphPad Prism 9.4.0.

