## Supplemental Figures for "A multiplexed, target-based phenotypic screening platform using CRISPR interference in *Mycobacterium abscessus*"

Supplementary Figure 1: Read count distribution in *M. abscessus* Tn-seq dataset.
Supplementary Figure 2: Two-plasmid transformation workflow for rapid hypomorph generation in *M. abscessus*.
Supplementary Figure 3: Verification of successful transformants using the two-plasmid methodology.
Supplementary Figure 4: Growth rate inhibition metrics across all strains in multiplex.
Supplementary Figure 5: Transcriptional responses to isoniazid treatment.

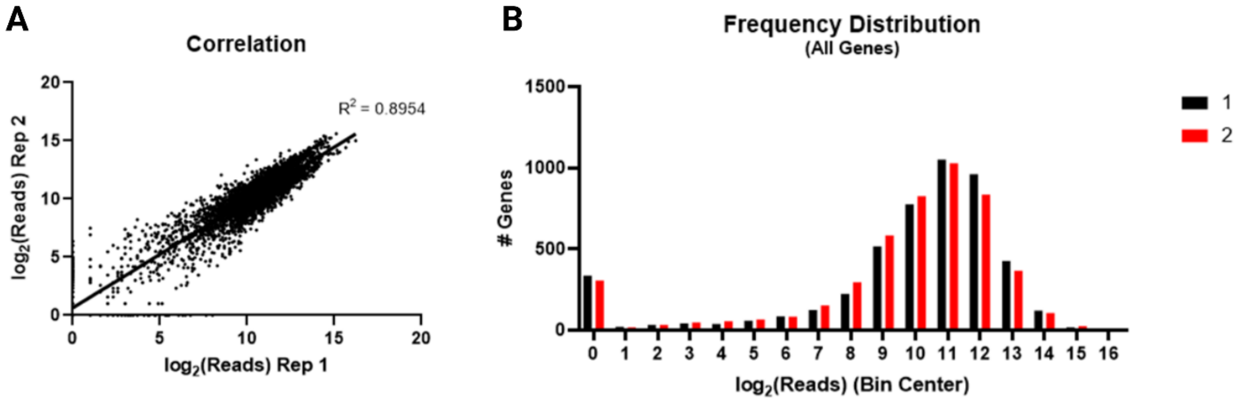


**Supplementary Figure 1: Read count distribution in *M. abscessus* Tn-seq dataset.**

**A.** Read count correlation between duplicate independent Tn-seq experiments. Each data point represents reads at one TA site. Pearson correlation $R^2 = 0.8954$.

**B.** Distribution of average read counts per gene for duplicate independent Tn-seq datasets. A bimodal distribution is observed with predicted essential genes on the left and predicted non-essential genes on the right.

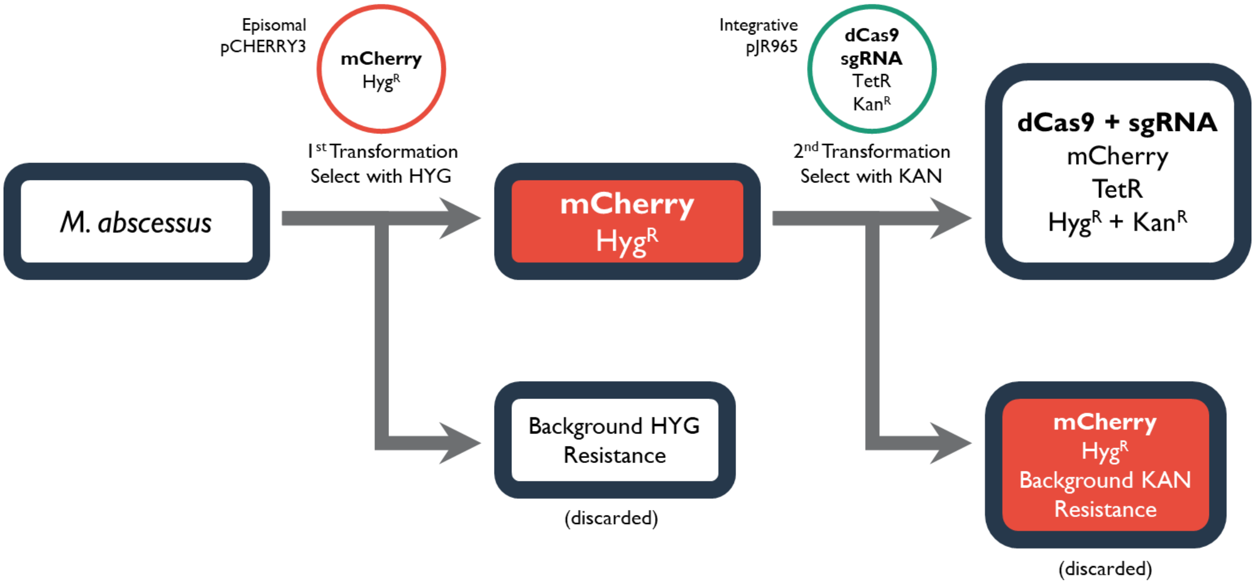


**Supplementary Figure 2: Two-plasmid transformation workflow for rapid hypomorph generation in *M. abscessus*.**

Briefly, WT *M. abscessus* is transformed with pCHERRY3 and selected with HYG. Successful transformants are identified as pink colonies and used for subsequent transformation with pJR965. After selection with KAN, successful transformants that integrated the CRISPRi machinery (as well as the TetR repressor) will then appear as white colonies, which can be selected from the pink background mutants that have acquired KAN resistance spontaneously.

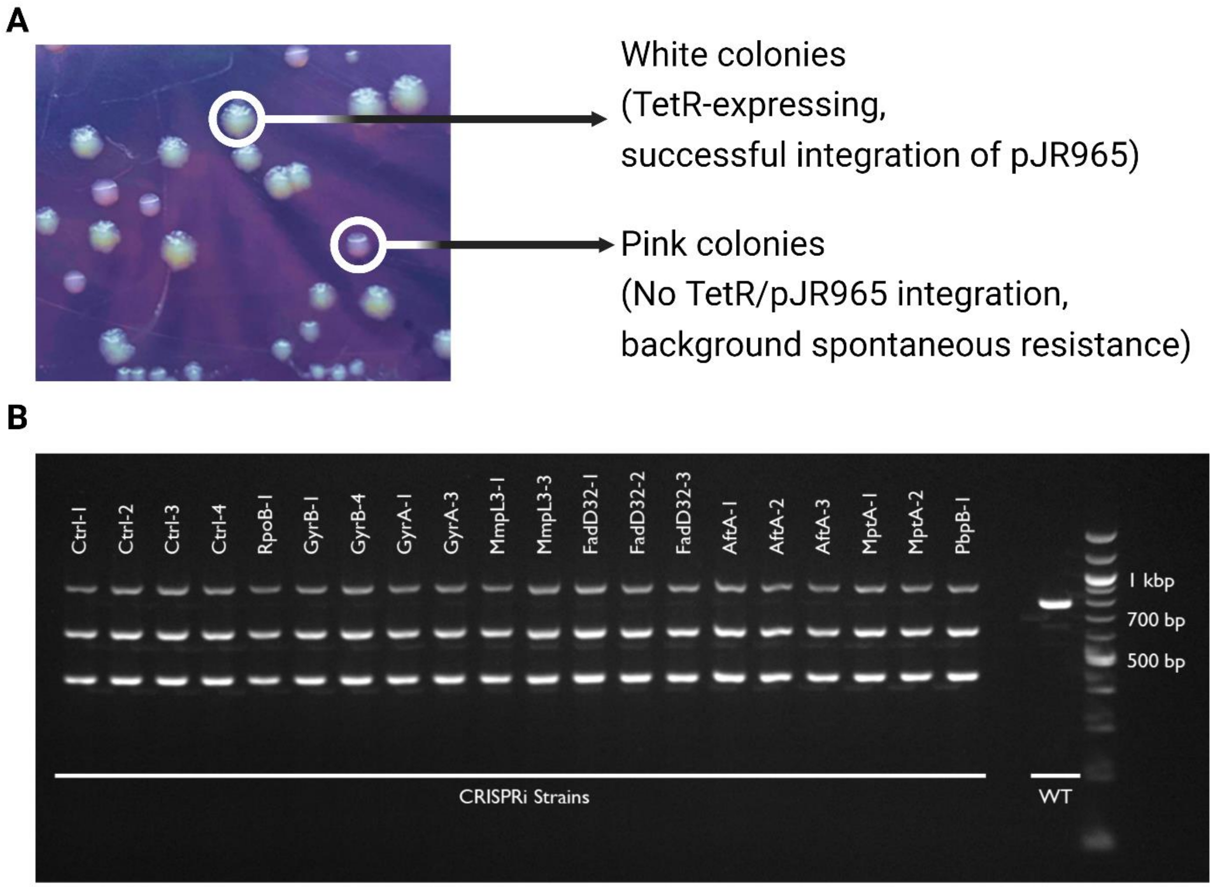


**Supplementary Figure 3: Verification of successful transformants using the two-plasmid methodology.**

**A.** White colonies can be easily identified visually after the second transformation with successful integration of the pJR965 plasmid leading to TetR expression which turns off mCherry expression. In contrast, background spontaneous resistance can be identified by the absence of pJR965 integration and TetR expression, which leaves mCherry expression on.

**B.** Representative gel images of colony PCR of CRISPRi hypomorph strains generated. In almost all cases, only a single colony was required to identify the correct transformants evidenced by the characteristic three bands (see Figure 2.3B, C) (labeled CRISPRi Strains below, with individual strain names listed above). A WT strain was included to portray band pattern for a background spontaneous resistant clone (second right-most lane). 1 kb ladder used in reference lane (right-most lane) with 500 bp, 700 bp and 1 kb bands labeled.

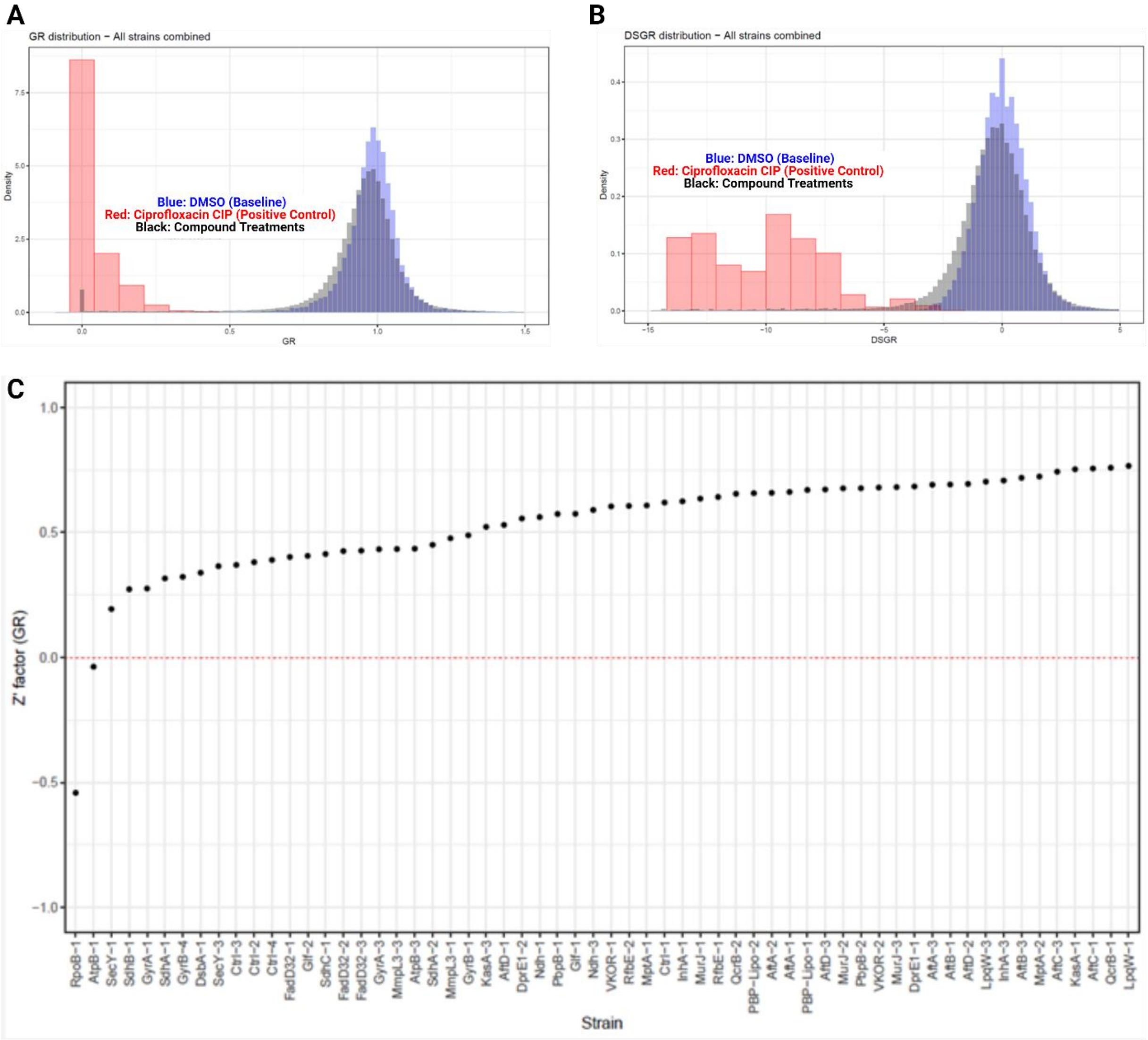


**Supplementary Figure 4: Growth rate inhibition metrics across all strains in multiplex.**

**A-B.** Distribution of GR (A) and DSGR (B) values across all strains. Values represent an average of duplicate treatments and are colored based on DMSO-treated wells (blue), CIP-treated wells (red), or compound treated-wells (grey/black).

**C.** *Z*' factors distributed across all 60 strains in multiplex. Z' factor was calculated using GR values in DMSO-treated vs. CIP-treated wells.

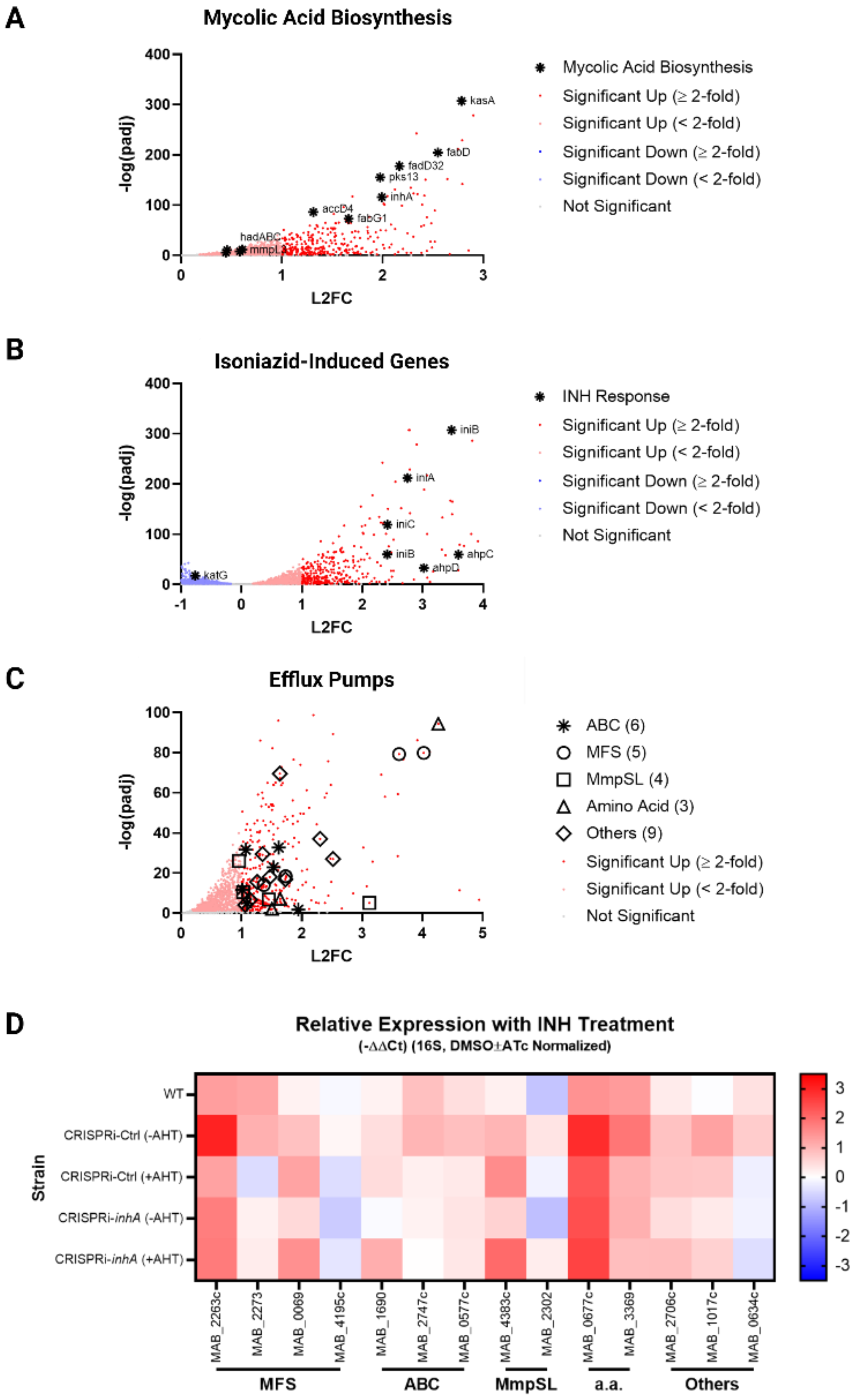


**Supplementary Figure 5: Transcriptional responses to isoniazid treatment.**

A-C. Differentially expressed genes from RNA-seq of isoniazid (INH)-treated WT *M. abscessus* compared to DMSO-treated bacteria. Genes with adjusted p-value < 0.05 were assigned as significantly differentially expressed. Genes involved in mycolic acid biosynthesis (A) and established isoniazid-induced genes (B) labeled with gene names. Efflux pumps from various families highlighted with black symbols (C).

D. qRT-PCR verification of a subset of predicted up-regulated efflux pumps in WT *M. abscessus*, or CRISPRi-Ctrl and CRISPRi-inhA with or without CRISPRi induction (± AHT). Relative expression calculated by normalizing Ct values to corresponding Ct values for 16S rRNA to determine ΔCt values, then normalized to matched ΔCt values for DMSO-treated samples (± AHT) to obtain ΔΔCt values. Data are mean values from triplicate samples plotted on a heat-map scaled from -3.5 (blue, significantly down-regulated) to 0 (white, no change) to 3.5 (red, significantly up-regulated).
